# PIP5k1 β controls bone homeostasis through modulating both osteoclast and osteoblast differentiation

**DOI:** 10.1101/288910

**Authors:** Xiaoying Zhao, Guoli Hu, Chuandong Wang, Lei Jiang, Jingyu Zhao, Jiake Xu, Xiaoling Zhang

## Abstract

PIP5K1β is crucial to generation of phosphotidylinosotol (4, 5) P_2_. PIP5K1β participates in numerous cellular activities, such as B cell and platelet activation, cell phagocytosis and endocytosis, cell apoptosis, and cytoskeletal organization. In the present work, we aimed to make insight into the function of PIP5K1β in osteoclastogenesis and osteogenesis to provide promising strategies for osteoporosis prevention and treatment. We discovered that PIP5k1β deletion in mice resulted in obvious bone loss and PIP5K1β was highly expressed both during osteoclast and osteoblast differentiation, besides, PIP5K1β deletion enhanced the proliferation and migration of BMMs to promote osteoclast differentiation. PIP5k1β^−/−^ osteoclast exhibited normal cytoskeleton architecture but stronger resorption activity. PIP5k1β deficiency also promoted activation of MAPK and Akt signaling, enhanced TRAF6 and c-Fos expression, facilitated the expression and nuclear translocation of NFATC1 and upregulated Grb2 expression, thereby accelerating osteoclast differentiation and function. Finally, PIP5K1β enhanced osteoblast differentiation by upregulating master genes expression through triggering smad1/5/8 signaling. Thereby, PIP5K1β modulate bone homeostasis and remodeling.

## Introduction

Bone remodeling depends on the dynamic balance and precise coordination of bone resorption and subsequent bone formation, which are driven by osteoclast and osteoblast activation, respectively. Disturbances of this delicate balance lead to skeletal diseases, such as osteopenia, osteoporosis and osteopetrosis (Boyle et al, 2003). Bone-resorbing osteoclasts are multinucleated cells derived from monocyte-macrophage precursors which originated from hematopoietic stem cells, whereas bone-forming osteoblasts are derived from mesenchymal stem cells (MSC)(Boyle et al, 2003; Dennis et al, 1999; Pittenger et al, 1999; Teitelbaum, 2000). Osteoblast lineage cells produce RANKL and stimulate their RANK receptors on osteoclast precursors, resulting in osteoclast differentiation by activating the downstream signaling pathways such as transcription factors NF-κB, c-Fos, and nuclear factor of activated T cells c1 (NFATc1)(Teitelbaum & Ross, 2003). To initiate bone resorption, osteoclasts need to successfully assemble a typical adhesion structure that is called actin ring or sealing zone, which consists of a belt of densely packed podosomes interlinked by an acto-myosin network (Georgess et al, 2014; Touaitahuata et al, 2014).

Exacerbated by general population aging, osteoporosis has become a dominating health problem around the world (Sambrook & Cooper, 2006). Various conditions can provoke osteoporosis, such as bone metastasis, disability and inflammation, furthermore, in postmenopausal women, estrogen deficiency gives rise to excessive bone loss.(Touaitahuata et al, 2016) Despite recent advances in bone biology, the precise molecular mechanisms responsible for pathological osteoporosis remain elusive. Therefore, clarifying the molecular mechanisms and novel molecules involved in the maintenance of bone homeostasis is critical for deeper understanding of skeletal health and development of novel therapeutics against various bone disorders.

Lipid kinases and their phosphoinositide products perform essential functions in secretory vesicle trafficking. Phosphoinositide (PI) contributes to numerous basic biological processes, such as chemotaxis, intercellular trafficking, polarity formation and cytokinesis. (Takenawa & Itoh, 2001) PIs are essential not only as membrane components in Eukaryotes and as precursors of second messengers like IP3 and PIP3 but also act as specialized membrane docking sites for effectors of diverse signaling cascades.(Takenawa & Itoh, 2001) Accumulating evidence indicates that PIs and PI-interacting proteins such as Rho, Arf, and Rab small GTPases serve as modulators of osteoclast differentiation.(Chellaiah, 2006; Ory et al, 2008) PIs, especially PIP2, PIP3 and IP3, have been reported to serve various essential roles during osteoclast differentiation and function as well as in maintain bone homeostasis. PIP5K1β is a member of the Type 1 phosphatidylinositol 4-phosphate 5-kinases (PIP5K1s; α, β, and γ), which are a family of isoenzymes producing phosphatidylinositol 4, 5-bisphosphate [PI (4, 5)P2] using phosphatidylinositol 4-phosphate as substrate(De Matteis & Godi, 2004; Ishihara et al, 1996; Ishihara et al, 1998; Kanaho et al, 2007; Oude Weernink et al, 2004; van den Bout & Divecha, 2009). PIP5K1s have been reported to be involved in a wide range of cellular functions not only as an enzyme generating PIP2, which is a critical regulator of cell adhesion formation, actin dynamics, and membrane trafficking, but also as cell signaling modulation factor, such as cytoskeleton assembly, exocytosis, endocytosis, cell apoptosis, and so on(Mao & Yin, 2007; van den Bout & Divecha, 2009). PIP5k1γ, which is a major PtdIns (4,5)P2-synthesising enzyme in the rodent brain with three splicing variants, is elegantly involved in neurons and neuroendocrine cells: Hara Y et al reported that PIP5K1γ, particularly PIP5K1γ_i2, performs essential roles in neuronal migration, possibly through recruitment of adhesion components such as talin and focal adhesion kinase to the plasma membrane.(Hara et al, 2013) In murine megakaryocytes, PIP5K1γ defect results in plasma membrane blebbing accompanied by a decreased connection of the membrane to the cytoskeleton possibly through a pathway involving talin.(Wang et al, 2008b) Moreover, PIP5K1γ_i2 revealed unique regulatory and targeting mechanisms through its C-terminal 26 amino acids. In neurons, PIP5K1γ_i2 modulates the clathrin-mediated endocytosis of synaptic vesicles at presynapses and α-amino-3-hydroxy-5-methyl-4-isoxazolepropionate-type glutamate receptors during long-term depression at postsynapses via association with adaptor protein complex (AP)-2(Unoki et al, 2012). In non-neuronal cells, PIP5K1γ_i2 also regulates the formation of focal adhesions through the interaction with talin.(Di Paolo et al, 2002; Ling et al, 2002; Sun et al, 2007) Furthermore, targeted disruption of PIP5K1γ leads to widespread developmental and cellular defects. PIP5K1γ-null embryos have myocardial developmental defects including impaired intracellular junctions resulting in heart failure and extensive lethality at embryonic day 11.5 as well as impaired PIP2 production, adhesion junction formation, and neuronal cell migration that lead to neural tube closure defects.(Wang et al, 2007) PIP5k1α, was reported to selectively modulate apical endocytosis in polarized renal epithelial cells and to regulate invadopodia formation and ECM degradation in human breast cancer cells by localized production of PI(4,5)P(2) (Szalinski et al, 2013). The function of PIP5k1β was investigated less and the roles of PIP5k1s in maintaining bone homeostasis were rarely reported. Only one report indicated that PIP5k1γ deficiency or overexpression delayed osteoclast differentiation and excess of PIP5k1γ disrupted osteoclasts cytoskeleton in a talin-independent way.(Zhu et al, 2013)

In the present study, we found that PIP5k1β was highly expressed during RANKL-induced osteoclast differentiation and PIP5k1β deletion in mice resulted in bone loss. To further understand the role of PIP5k1β in bone homeostasis, we investigated the functions of PIP5k1β acting on osteoclast and osteoblast differentiation by gain- and loss-of-function in vitro. We found that PIP5K1β can repress the proliferation and migration of bone-marrow–derived macrophage-like cells (BMMs) to inhibit osteoclast differentiation. Furthermore, PIP5k1β deletion promoted the activation of MAPK and Akt signaling cascades and enhanced expression of TRAF6 and c-Fos to facilitate NFATC1 expression and nuclear translocation, thereby accelerating osteoclast differentiation and function. Last but not the least, PIP5k1β deficiency upregulated Grb2 expression during osteoclast differentiation. PIP5K1β also enhanced osteoblast differentiation through activating of smad1/5/8. Thus, PIP5K1β can regulate bone mass and bone remodeling.

## Results

### PIP5k1β-deficient mice show an osteoporosis bone phenotype

Having demonstrated that PIP5k1γ served an essential role in modulating osteoclast differentiation and that it must be expressed at an actually exact level to maintain normal differentiation of osteoclast.(Zhu et al, 2013) To determine the role of PIP5k1β in the skeleton, we evaluated the mutant mouse strain with piggyback (PB) transposition system-induced mutation in the *PIP5k1β* gene. Homozygous mutation of PIP5k1β were born healthy at the predicted Mendelian frequencies. PIP5k1β^−/−^ mice had a similar body structure to that of WT littermates with no obvious differences observed between PIP5k1β^−/−^ and WT controls (Supplementary Fig. 1A). Interestingly, micro computed tomography (micro CT) analysis revealed that the trabecular bone mass density (BMD), trabecular bone volume versus tissue volume (BV/TV), trabecular bone number (Tb.N.) and trabecular bone thickness (Tb.Th.) were significantly lower but trabecular bone separation (Tb.Sp.) is obvious high in PIP5k1β^−/−^ mice compared to that of age- and sex-matched WT littermates (Fig. 1A–F), suggesting defects in trabecular bone growth. To gain further insight into the *in vivo* cellular phenotype of the PIP5k1β^−/−^ mice, bone histomorphometry was performed on decalcified sections stained for TRAP activity and for hematoxylin and eosin staining (Fig. 1G). Consistent with micro CT data, histomorphometric analysis of tibia from 8-week-old PIP5k1β^−/−^ mice revealed a lower trabecular bone mass when compared to WT mice (Fig. 1G, H). Analysis of osteoclast parameters using TRAP stained sections showed that PIP5k1β^−/−^ mice exhibited a significant increase in the number of osteoclasts (Fig. 1G, I). Moreover, Dynamic histomorphometric analysis detected the reduced bone formation rate and mineral apposition rate in PIP5k1β^−/−^ mice (Fig. 1J, K). Furthermore, the serum levels of CTX-1, a marker for bone resorption, was significantly increased (Fig. 1L) but serum OCN levels, a marker for bone formation, was dramatically decreased (Fig. 1O) in PIP5k1β^−/−^ mice. Moreover, immunohistochemistry detected that expression of CTSK, one of major markers of mature osteoclasts, in PIP5k1β deletion mice was significantly higher than that in wide type mice (Fig. 1M-N). These accumulating results suggest that PIP5k1β deletion leads to significant decrease in bone mass, which is possibly due to imbalance changes in osteoclasts bone resorption and osteoblasts bone formation.

**Figure 1.**
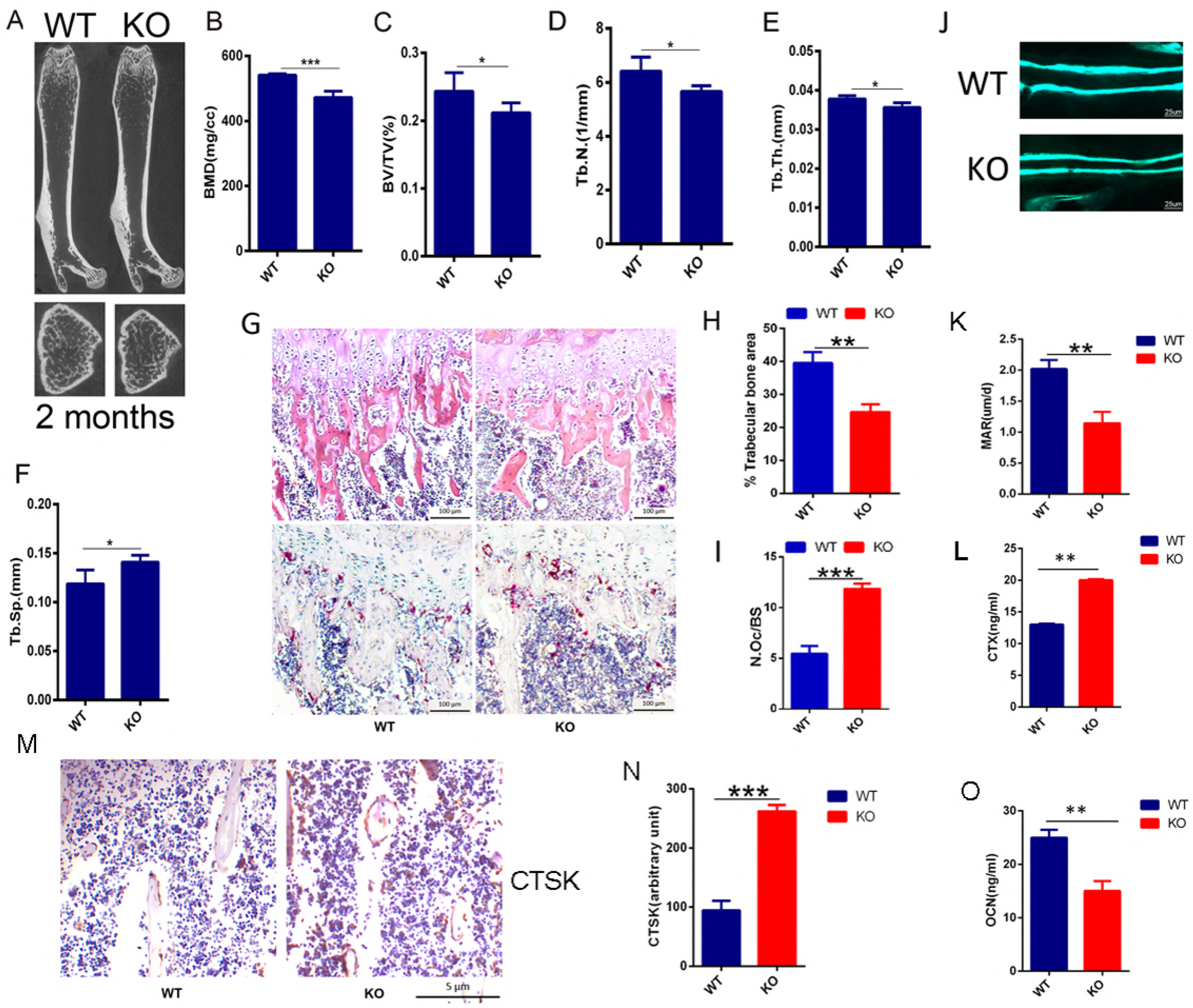
PIP5k1β-deficient mice exhibit an osteoporotic bone phenotype. (A) Representative micro-CT images of femurs in WT or PIP5k1β knockout 2-month-old male mice. (B-F) Bone mineral density (BMD), Bone volume per tissue volume (BV/TV), trabecular number (Tb.N.), trabecular bone thickness (Tb.Th.) and trabecular separation (Tb.Sp.) were assessed from the micro-CT measurements (n = 10). (G) The hematoxylin/eosin (H&E) (up) and TRAP staining (down) of histological section of proximal tibiae. Bars, 100 µm. (n = 10). (H-I) Analysis of trabecular bone area and quantification of N.Oc/BS (osteoclast number/ bone surface) in G. (J-K) Representative images of new bone formation (J) and quantification of MAR (K) as assayed by calcein double labeling. (n = 6). (L) Serum concentration of CTx-I in WT or PIP5k1β knockout mice (n = 6). (M-N) The expression of CTSK in WT or PIP5k1β knockout mice shown by immunohistochemistry (M) and the staining density were quantified by ImageJ (N). (n=6 per group) (O) Serum concentration of OCN in WT or PIP5k1β knockout mice (n = 6) (Data represent as means ± S.D. of three independent experiments. *P < 0.05; **P < 0.01; ***P < 0.001.)

### PIP5k1β expression was increased during RANKL-induced osteoclast differentiation

Thereby, to determine how PIP5k1β influences osteoclasts formation and function, we first examined the transcriptomes of preosteoclasts and mature osteoclasts differentiated from mouse bone marrow–derived macrophage-like cells (BMMs) using cDNA microarray and found that among PIP5k1s only PIP5k1β was highly expressed in mature osteoclasts.(Fig. 2A). When we cultured BMMs in the presence of RANKL and M-CSF, the expression of PIP5k1β was markedly increased during RANKL-induced osteoclast differentiation since the third day of RANKL induction, however, the expressions of PIP5k1α and PIP5k1γ were not changed (Fig. 2B, Fig.3B). Expression levels of VATPased2 and TRAP, important markers of osteoclastogenesis, increased as the cells differentiated (Fig. 2B). To confirm the importance of PIP5k1β in skeletal biology, we explored the expression pattern of PIP5k1β in different mice organs such as bone, spleen, brain, thymus, lung, kidney, liver, heart, and muscle and found that PIP5k1β was highly expressed in heart, lung, brain and bone (Supplementary Fig. 1B), which indicated that PIP5k1β might play a crucial role in bone biology. These results indicate that PIP5k1β might be involved in regulation of osteoclastogenesis and bone remodeling.

**Figure 2.**
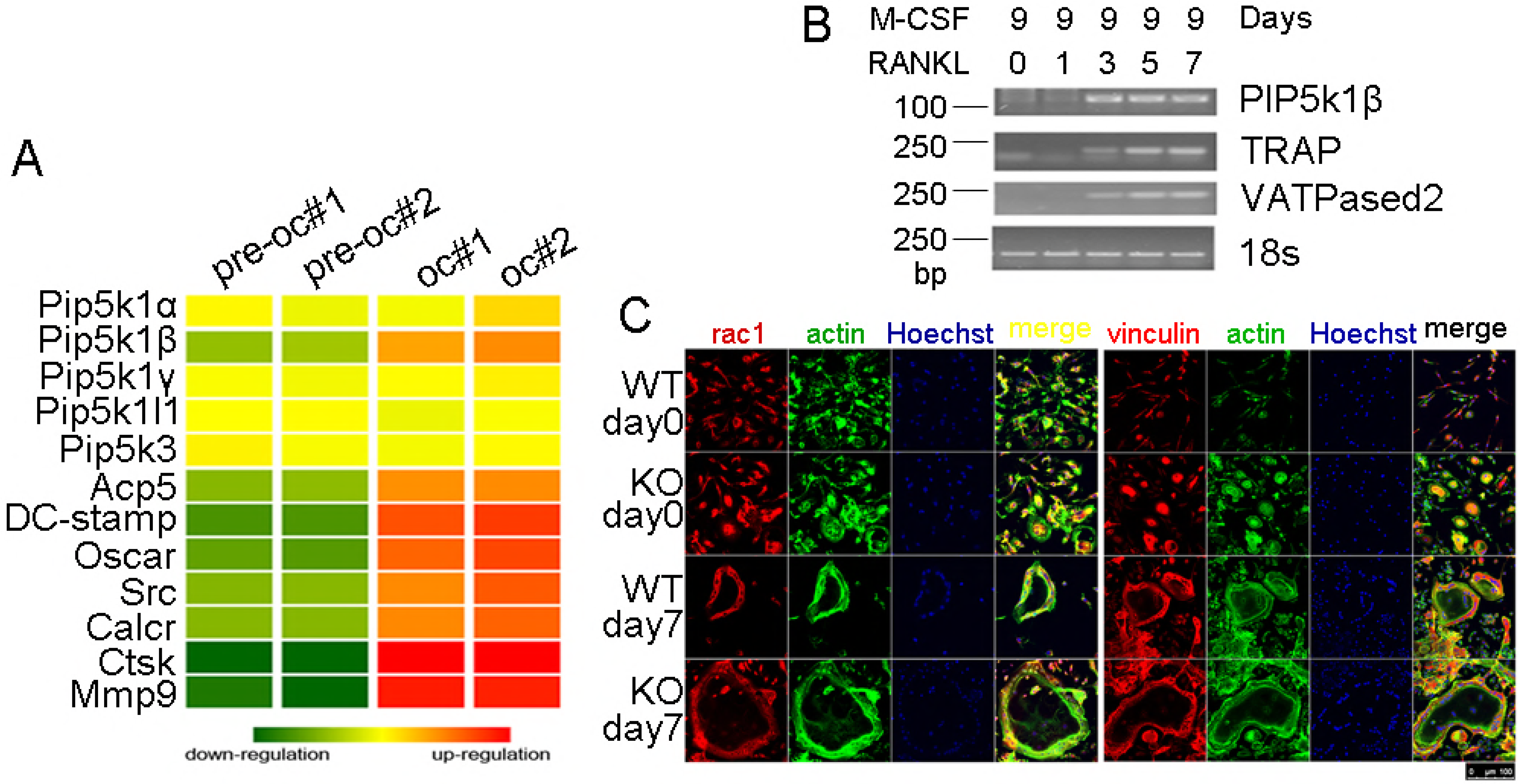
PIP5k1β expression was increased during RANKL-induced osteoclast differentiation and PIP5k1β-deficient osteoclasts exhibit normal morphology. (A) cDNA from preosteoclasts and mature osteoclasts were subjected to Micro array assay and the expression levels of PIP5k family genes and osteoclasts master genes were detected. The heat map is ordered by degree of differential expression of the indicated genes between preosteoclasts and mature osteoclasts. (B) RT-PCR detection for the expression levels of PIP5k1β, TRAP, VATPased2 during osteoclastogenesis and 18s RNA was used as control. (C) Bone marrow cells from WT or PIP5k1β−/− mice were treated by 15 ng/ml M-CSF for two days and the cells adhesion to the cell plates (named BMMs) were underwent osteoclastogenesis by stimulation with 20 ng/ml M-CSF and 75 ng/ml RANKL for 7 days. The actin ring formation, Vinculin, and Rac1 expression and colocalization with F-actin were assessed by classical confocal microscopy. F-actin was stained with phalloidin (green) and nuclear was stained with Hoechst 3342. (All the data were confirmed by three independent experiments.)

**Figure 3.**
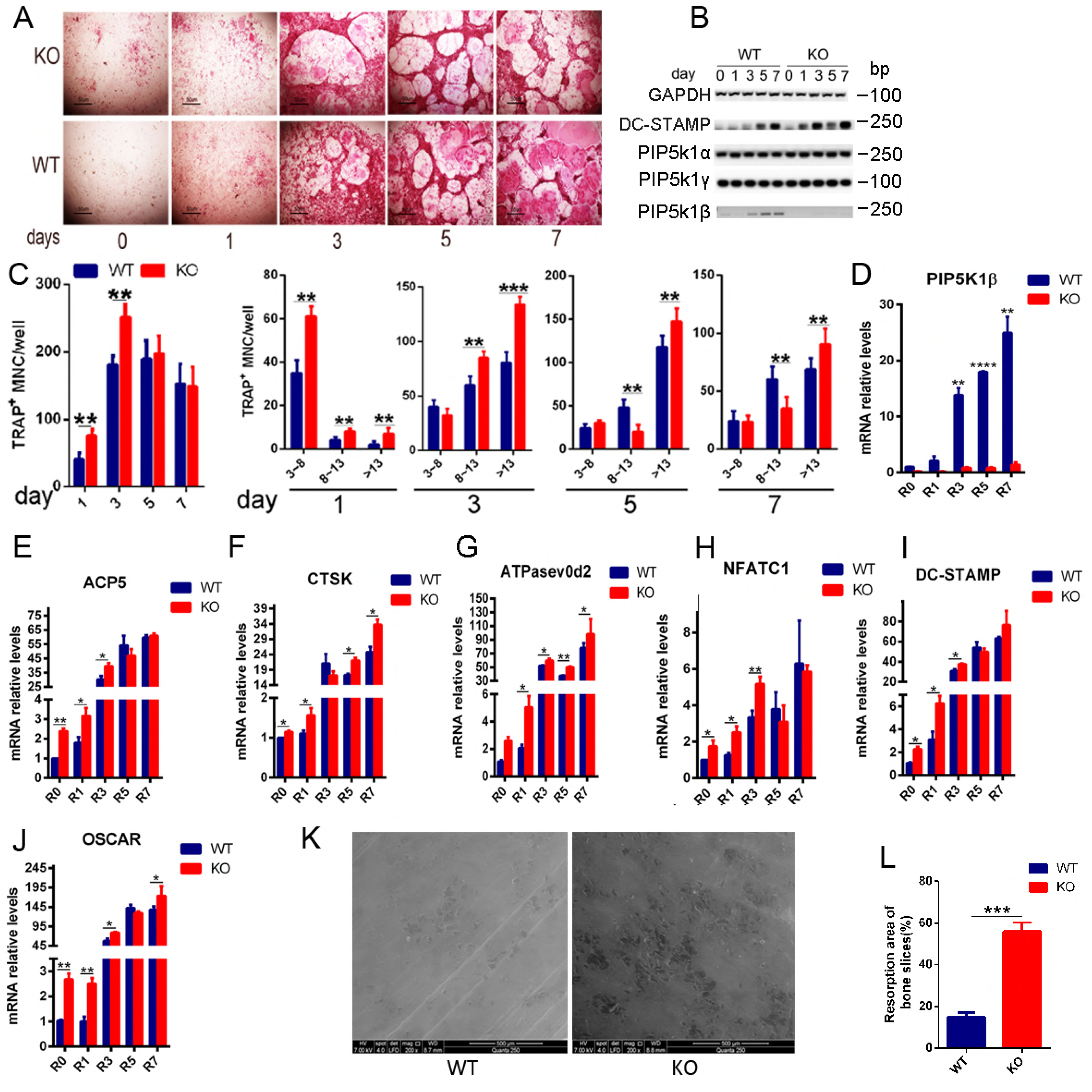
Deficiency of PIP5k1β enhanced osteoclast differentiation and bone resorption. Bone marrow cells from WT or PIP5k1β−/− mice were treated by 15 ng/ml M-CSF for two days and the cells adhesion to the cell plates underwent osteoclastogenesis by stimulation with 20 ng/ml M-CSF and 75 ng/ml RANKL for the indicated number of days, the osteoclasts formation were detected by TRAP staining (A) and the osteoclasts number were counted (C) (n=3). (B, D-J) WT or PIP5k1β−/− BMMs were cultured with M-CSF (20 ng/mL) and RANKL (75 ng/mL) for 0, 1, 3, 5 or 7 days. PIP5k1β, PIP5k1α, PIP5k1γ and osteoclast-specific gene expression (ACP5, Ctsk, ATPasev0d2, NFATc1, DC-STAMP and OSCAR) were analyzed by RT-PCR (B) and real-time polymerase chain reaction (PCR) and the results were normalized to the expression of GAPDH (D-J). All experiments were performed at least three times. (K) WT (K left) or PIP5k1β−/− (K right) BMMs were cultured on bovine bone slices with 20 ng/ml M-CSF and 75 ng/ml RANKL for 8 days and the pit formation was detected by SEM and the pit formation rate was measured by Image J and presented graphically (L) (All the experiments were repeated three times. Data represent as means ± SD of three independent tests. *P< 0.05; **P < 0.01; ***P < 0.001 versus WT).

### PIP5k1β-deficient osteoclasts exhibit normal morphology

Accumulating evidences demonstrated that PIP5k1 family kinase serve central roles in cytoskeleton assembly(Kisseleva et al, 2005; Rozelle et al, 2000; Shibasaki et al, 1997; van den Bout & Divecha, 2009; van Horck et al, 2002; Wang et al, 2008b), so we give rise to the hypothesis that PIP5k1β deficient may affect osteoclast cytoskeleton. To verify this hypothesis, we detect the actin ring formation as well as the localization of vinculin and Rac1, which are critical factors that can contribute to the ability of osteoclasts to rearrange podosomes into the sealing zone and establish the bone-resorbing apparatus, through classical confocal microscopy(Croke et al, 2011; Fukunaga et al, 2014; Ory et al, 2000; Sun et al, 2005; Wang et al, 2008a). F-actin exhibits a similar well-defined peripheral belt architecture in both WT and PIP5k1β-deficient osteoclast, and PIP5k1β deletion promotes podosome formation in preosteoclasts and PIP5k1β deletion osteoclasts display much larger actin rings compared with WT osteoclasts. Vinculin and Rac1 localization and expression are also normal in PIP5k1β-deletion osteoclasts, which are all enriched at the podosome belt and are colocalized with F-actin (Touaitahuata et al, 2016) (Figure. 2C). Finally, the expression levels of β3 integrin, which involves in the association of osteoclasts with bone matrix to trigger osteoclast bone resorption activity, was also uninfluenced by PIP5k1β deficiency (Supplementary Fig. 1C, D).These results indicate that PIP5k1β deletion accelerates actin ring formation but does not affect osteoclast cytoskeleton assembly and sealing zone formation as well as interaction ability with bone matrix on the whole.

### Deficiency of PIP5k1β enhanced osteoclast differentiation and bone resorption

Given our observation that PIP5k1β expression is upregulated during RANKL-induced oseoclastogenesis, we examined the effect of PIP5k1β on osteoclast differentiation. Bone marrow cells from 9-week-old WT and PIP5k1β^−/−^ mice were cultured with M-CSF to generate WT and PIP5k1β^−/−^ BMMs, which were used as osteoclast precursor cells to be stimulated with RANKL and M-CSF. Deletion of PIP5k1β in BMMs significantly stimulates the formation of TRAP-positive multinucleated cells (TRAP-positive MNCs) mediated by RANKL (Fig. 3A, C). More osteoclasts with 3–8 nuclei or >13 nuclei were observed in the PIP5k1β-deficient group. Besides, RT-PCR results exhibited an increased expression of DC-STAMP in PIP5k1β^−/−^ BMMs compared with WT BMMs during RANKL-induced osteoclast differentiation (Fig. 3B). Consistently, gene expression analysis revealed that PIP5k1β expression was significantly increased since the third day of RANKL-induced osteoclastogenesis, but the expressions of the other two PIP5k1 family proteins PIP5k1α and PIP5k1γ were not obviously changed during both WT and PIP5k1β deletion BMMs differentiation into osteoclasts (Fig. 3B, D). Moreover, the expressions of the marker genes of osteoclast formation, such as TRAP, CTSK, ATPasev0d2, NFATC1, DC-STAMP and OSCAR were dramatically enhanced during PIP5k1β deficiency BMMs differentiation into osteoclasts compared with that of WT BMMs. (Fig. 3E-J) Furthermore, we cultured WT and PIP5k1β^−/−^ BMMs on bovine bone and initiated osteoclastogenensis by M-CSF and RANKL stimulation, respectively. After successful formation of mature osteoclasts, the cells were cultured for another day with M-CSF and RANKL stimulation to further resorb bone; the resorbed bones then underwent SEM. We discovered that PIP5k1β deletion promoted osteoclasts bone resorption and the resorption area of the bone slices by PIP5k1β^−/−^ osteoclasts was significantly larger than that of WT osteoclasts. (Fig. 3 K-L) These results suggest that PIP5k1β deficiency enhances osteoclast differentiation and formation as well as promotes osteoclasts bone resorption function; and this function might not depend on the compensational role of PIP5k1α and PIP5k1γ.

### Overexpression of PIP5k1β arrests osteoclastogenesis

To further verify the role of PIP5k1β in osteoclast differentiation and formation, we overexpress PIP5k1β in BMMs to evaluate its effect on RANKL-induced osteoclast differentiation. PIP5k1β was overexpressed in the BMMs by lentivirus transfection into the BMM cells and then the BMMs were cultured with RANKL and M-CSF. The overexpression levels of PIP5k1β with gradually increased dose of lentivirus were showed by RT-PCR (Fig. 4A). TRAP staining analysis showed that overexpression of PIP5k1β significantly inhibits osteoclast differentiation and the inhibition effect was gradually increased along with the increasing dose of PIP5k1β overexpression. (Fig. 4 B-E)

**Figure 4.**
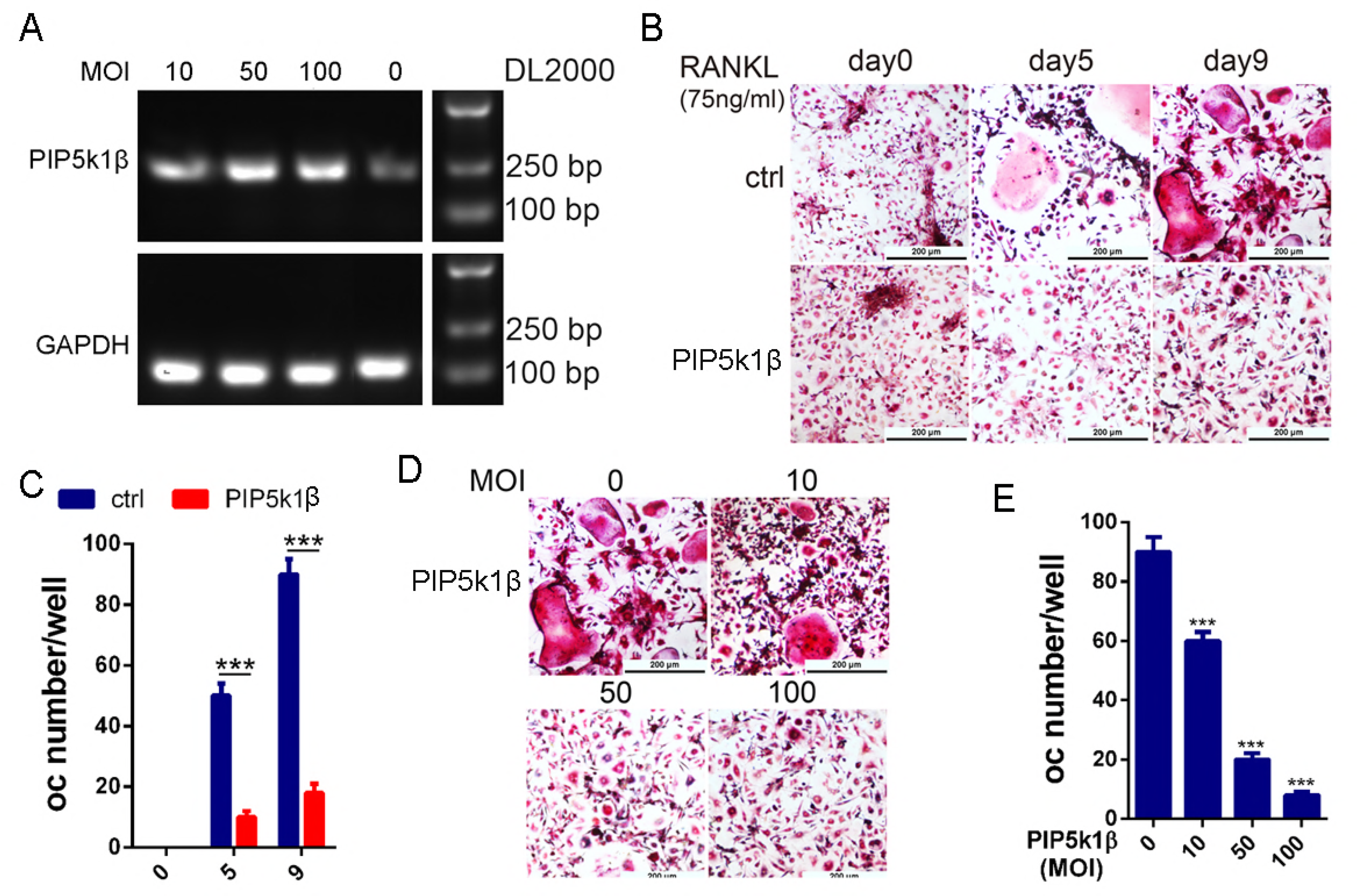
Overexpression of PIP5k1β arrests osteoclast formation. (A) WT BMMs were transduced with LV5-NC or gradually increased concentration of LV5-PIP5k1β, and the expression of PIP5K1β was detected by RT-PCR. (B-C) BMMs transduced with LV5-NC or 50 MOI of LV5-PIP5k1β were cultured with RANKL and M-CSF for 0, 5 or 9 days. The cells were stained for TRAP activity and osteoclasts number were counted. (D-E) BMMs from A were cultured with RANKL and M-CSF for 9 days. The cells were stained for TRAP activity and osteoclasts number were counted. (Scale bar, 200 μm, all the experiments were repeated three times. Data represent as means ± SD of three independent tests. ***, P < 0.001.)

### Deficiency of PIP5k1β enhances proliferation, migration and NFATc1 nuclear localization in preosteoclasts

To explore the mechanisms of PIP5k1β inhibition of osteoclast differentiation, we examined the effect of PIP5k1β on BMMs proliferation and migration. The wound healing assay displayed that PIP5k1β deletion promotes preosteoclasts migration triggered not only by M-CSF or by both M-CSF and RANKL (Fig. 5A, B). The CCK8 assay showed that PIP5k1β deficiency enhanced BMMs proliferation under the induction of RANKL and M-CSF, whereas only RANKL or M-CSF had no significant effect on the proliferation of BMMs (Fig. 5C). Furthermore, PIP5k1β deficiency facilitated NFATC1 expression and nuclear translocalization when triggered by RANKL detected by classical confocal microscopy and western blotting (Fig. 5D-E). Besides, luciferase assay with a NFATC1 responsive reporter was strongly dampened by PIP5k1β overexpression under the stimulation of RANKL (Fig. 5F-G). These results implied that PIP5k1β suppressed osteoclast differentiation by depressing preosteoclast proliferation, migration and also reducing NFATC1 signaling by decreasing its expression and nuclear translocalization.

**Figure 5.**
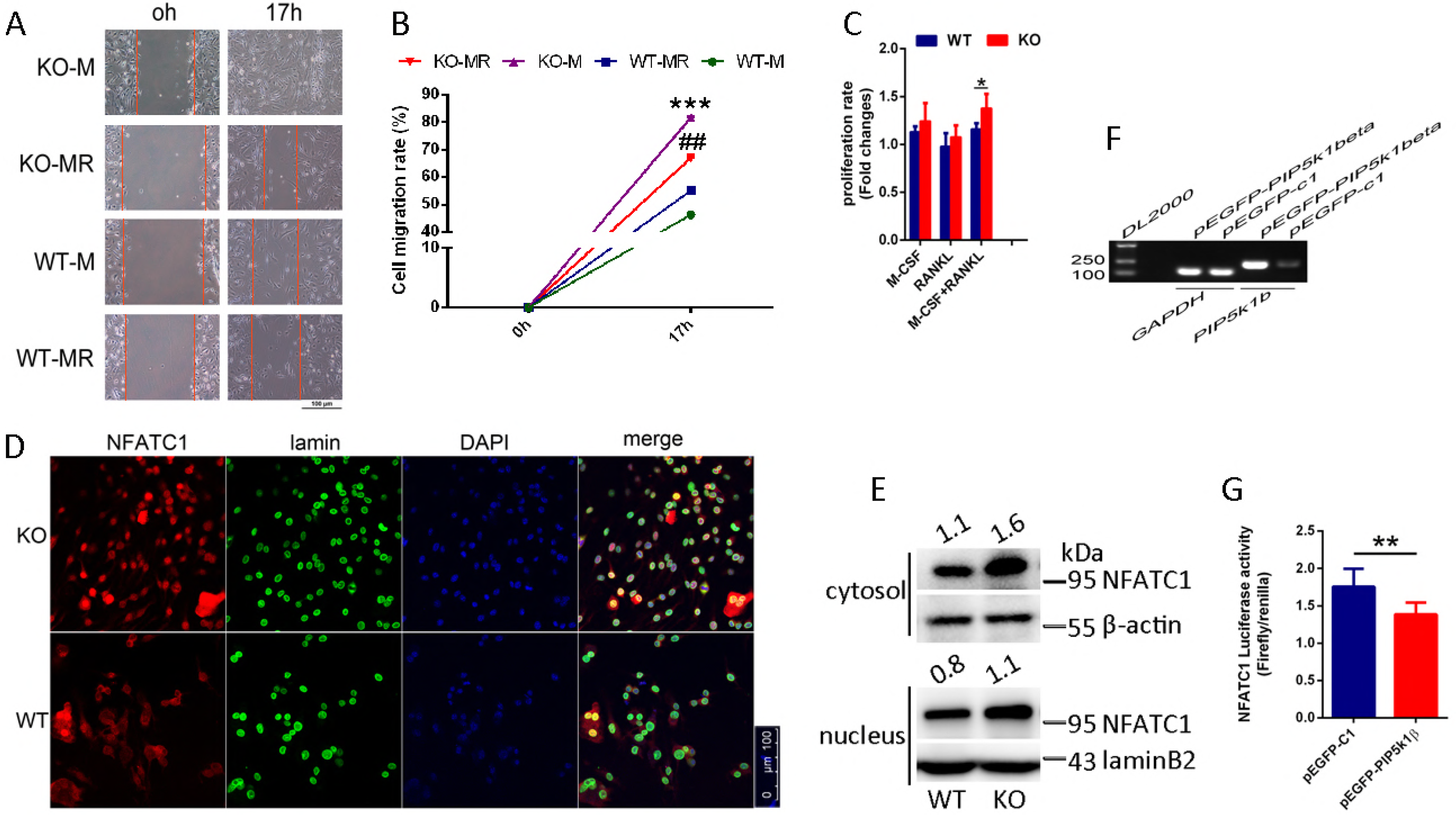
Deficiency of PIP5k1β enhances proliferation, migration and NFATc1 nuclear localization in preosteoclasts. (A) The wound healing assay. WT or PIP5k1β−/− BMMs were cultured in 48-well plates and serum- and cytokine-starved for 12 h; scratches were made and serum-free culture medium with M-CSF or both M-CSF and RANKL was added. Cells migrating to the scratches were monitored and images were taken at 0 and 17 h after wounding. (B) The migration rate was calculated according to the equation: percentage wound healing = ((wound length at 0 h) − (wound length at 17 h))/(wound length at 0 h) ×100 (C) CCK8 assay was performed to text the proliferation rate of WT or PIP5k1β−/− BMMs under triggering with M-CSF, RANKL, or both. (D-E) Nuclear translocation of NFATC1 in WT or PIP5k1β−/− osteoclasts stimulating with M-CSF and RANKL was detected by immunofluorescence (D) and western blotting (E). (F) RAW264.7 cells stably transfected with an NFATC1 luciferase reporter construct were seeded in 48-well plates and maintained in the cell culture medium for 24 h. The cells were then transduced with vector or pEGFP-PIP5k1β for 24 h, followed by incubation with RANKL (100 ng/mL) for 4 h. NFATC1 luciferase activity was measured (down) and PIP5k1β mRNA levels were detected via RT-PCR (up). (All experiments were repeated three times. Data represent as means ± SD of triplicate tests. *P< 0.01; **, P < 0.001; ## P< 0.01; ***, P < 0.0001 versus WT).

### Deficiency of PIP5k1β leads to increases of M-CSF- and RANKL-mediated MAPK, c-Fos and AKT signaling cascades

PIP5k1β was discovered to be highly expressed during RANKL-mediated osteoclastogenesis and PIP5k1β suppressed osteoclast formation and bone resorption capacity. We asked which signaling pathway modulated osteoclast differentiation and whose function is affected by the absence of PIP5K1β, which in turn affected osteoclastogenesis. First, we examined the archetypical RANKL signaling mediators such as c-Fos, TRAF6, NFATC1, Cathepsin K (CTSK) and TRAP, and found that deletion of PIP5k1β prominently enhanced the expression of these genes during RANKL-mediated osteoclstogenesis (Fig. 3E-F, H, Fig. 6A-D). During RANKL-induced osteoclastogenesis, ligation of RANKL with its receptors RANK leads to recruitment of TRAF6 by RANK, which leads to activation of nuclear factor-КB (NF-КB), the mitogen-activated kinases (MAPKs) and the Akt pathways.(Kadono et al, 2005; Kobayashi et al, 2001) In this context, we detected which of these pathways were modulated by PIP5k1β. The RANKL-induced activating phosphorylation of Akt and three typical MAP kinases (p38, ERK1/2 and JNK) were substantially elevated in PIP5k1β deletion BMMs compared with that of WT BMMs (Fig. 6E-J).

**Figure 6.**
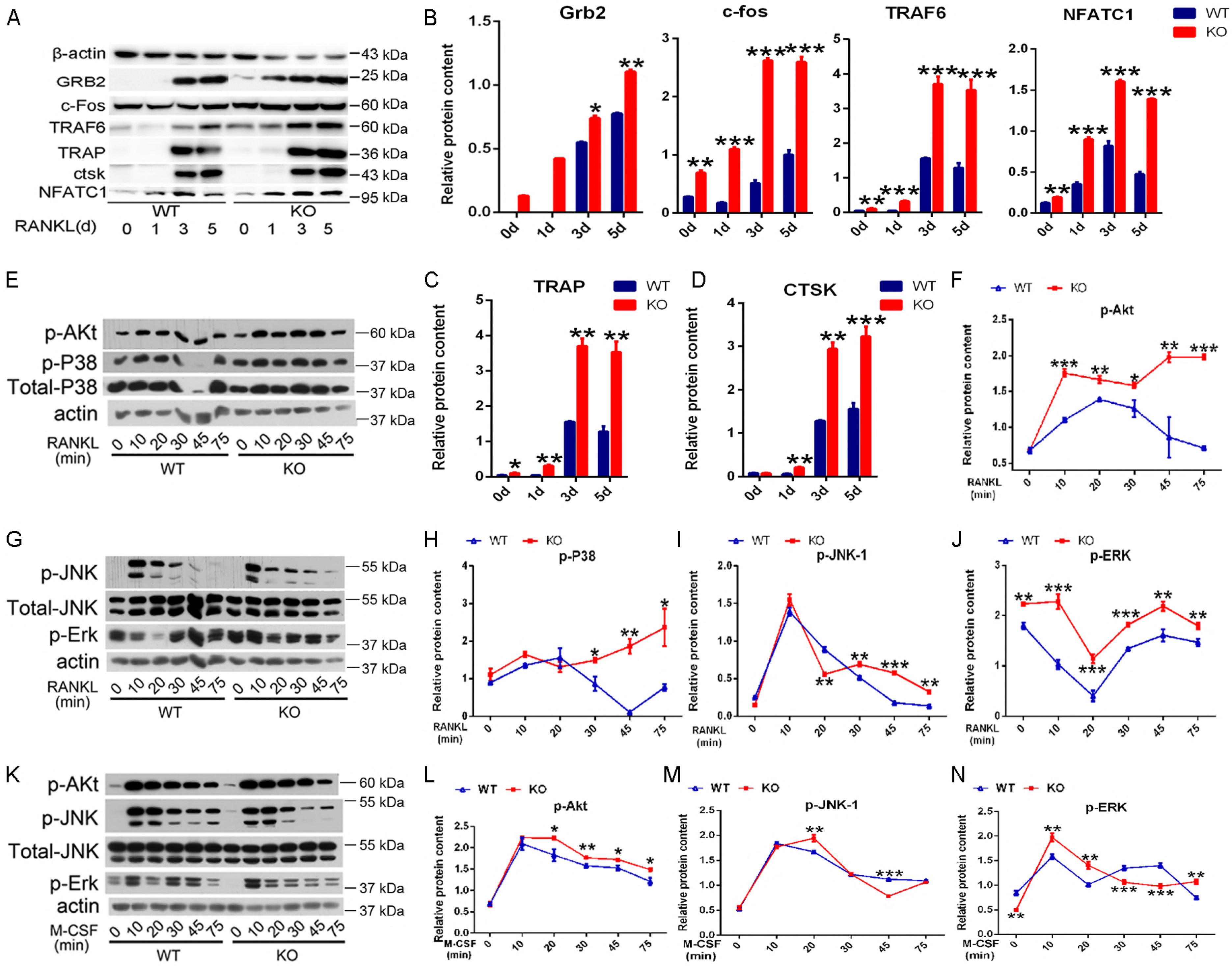
Deficiency of PIP5k1β leads to increases of M-CSF- and RANKL-mediated MAPK, c-Fos and AKT signaling cascades. (A-D) WT or PIP5k1β^−/−^ BMMs were cultured with M-CSF (20 ng/mL) and RANKL (75 ng/mL) for 0, 1, 3, 5days. Expression of TRAF6, c-Fos, Grb2, TRAP and Ctsk were analyzed by western blotting (A), the band densitometry were quantified and normalized to β-actin using ImageJ and graphically (B-D). Actin served as loading controls. (E-J) Serum- and cytokine-starved WT and PIP5KIβ^−/−^ macrophages were exposed to RANKL (100 ng/ml), Phosphorylated AKT, JNK, ERK, p38 were immunoblotted with time. Actin served as loading controls, and the expression levels were analyzed by ImageJ. (K-N) Serum- and cytokine-starved WT and PIP5KIβ^−/−^ macrophages were exposed to M-CSF (30 ng/ml). Phosphorylated AKT, ERK, JNK were immunoblotted with time. Actin served as loading control, and the expression levels were analyzed by ImageJ. (All experiments were repeated three times. Data represent means ± SD. *P< 0.05; **P< 0.01; ***, P < 0.001 versus WT).

Then, we treated WT and PIP5KIβ^−/−^ mice osteoclast precursors with M-CSF and assessed activation of established effector molecules, which mediated osteoclast formation. Absence of PIP5K1β had certain effects on M-CSF-induced phosphorylation of ERK, JNK or AKT in triggering with M-CSF but the enhance effects were not consistent along with the time of cells exposed to M-CSF, especially for JNK and ERK1/2, compared with WT BMMs (Fig.6K-N). Notably, the expressions of Grb2 was also enhanced during RANKL-mediated osteoclastogenesis in PIP5k1β^−/−^ osteoclasts compared with WT osteoclasts (Fig.6A). The adaptor protein Grb2 has been reported to play a pivotal role in several tyrosine kinase signal transduction pathways and previous study demonstrated that Grb2 promotes osteoclasts survival through activating Erk activation and bone resorption activity by enhancing their adhesion.(Levy-Apter et al, 2014) These data suggested that PIP5k1β deficiency influenced M-CSF or RANKL-induced osteoclastogenic properties to a certain extent, particularly RANKL-induced c-Fos, MAPK and AKT signaling pathways. Taken together, PIP5k1β suppressed osteoclasts differentiation and function might through influencing M-CSF- and RANKL-mediated MAPK, c-Fos and AKT signaling cascades.

### PIP5k1β accelerates osteoblast differentiation

Given that the serum bone formation marker OCN was significantly decreased in PIP5k1β^−/−^ mice compared with WT mice, we explored the effect of PIP5k1β on osteoblast differentiation and function to deepen our understanding of the role of PIP5k1β in bone homeostasis. First, immunohistochemistry revealed that expression of OSX, one of major transcriptional factors that control osteoblast differentiation, in PIP5k1β deletion mice was notably lower than that in wide type mice (Fig. 7A-B). Then, BMSCs constantly overexpress or knockdown of PIP5k1β were obtained by corresponding lentivirus transfection (Fig. 7C) and underwent osteoblast differentiation by stimulation with osteogenic medium. MTT assay revealed that knockdown or overexpression of PIP5k1β have no effect on BMSCs proliferation (Supplementary Fig.2A). Real-time PCR was performed to determine whether osteoblast marker genes were differentially expressed between WT and PIP5k1β overexpression osteoblasts. The mRNA levels of PIP5k1β were significantly enhanced during osteoblast differentiation and the mRNA expression of osteoblast marker genes, ALP, SP7, colla1, OCN and BSP, were increased by overexpression of PIP5k1β (Fig. 7D-K). Increased ALP activity was observed in PIP5k1β-overexpression osteoblasts at day 7 of culture and decreased ALP activity was observed in PIP5k1β-knockdown osteoblasts at day 7 of culture as compared to WT osteoblasts by ALP staining (Fig. 7L). The bone mineralization activity of osteoblasts derived by BMSCs constantly overexpression or knockdown of PIP5k1β was also altered consistently with ALP activity alteration relative to WT osteoblasts, as determined by alizarin red S staining and analysis at day 14 of culture (Fig. 7M). Furthermore, the protein levels of Col1α1, RUNX2 and OSX, which were central marker genes of osteoblastogenesis were all enhanced via PIP5k1β overexpression and decreased when PIP5k1β was knockdown. (Fig. 7N, supplementary Fig. 2B-C). These results implied that PIP5k1β enhanced MSCs differentiation into osteoblast.

**Figure 7.**
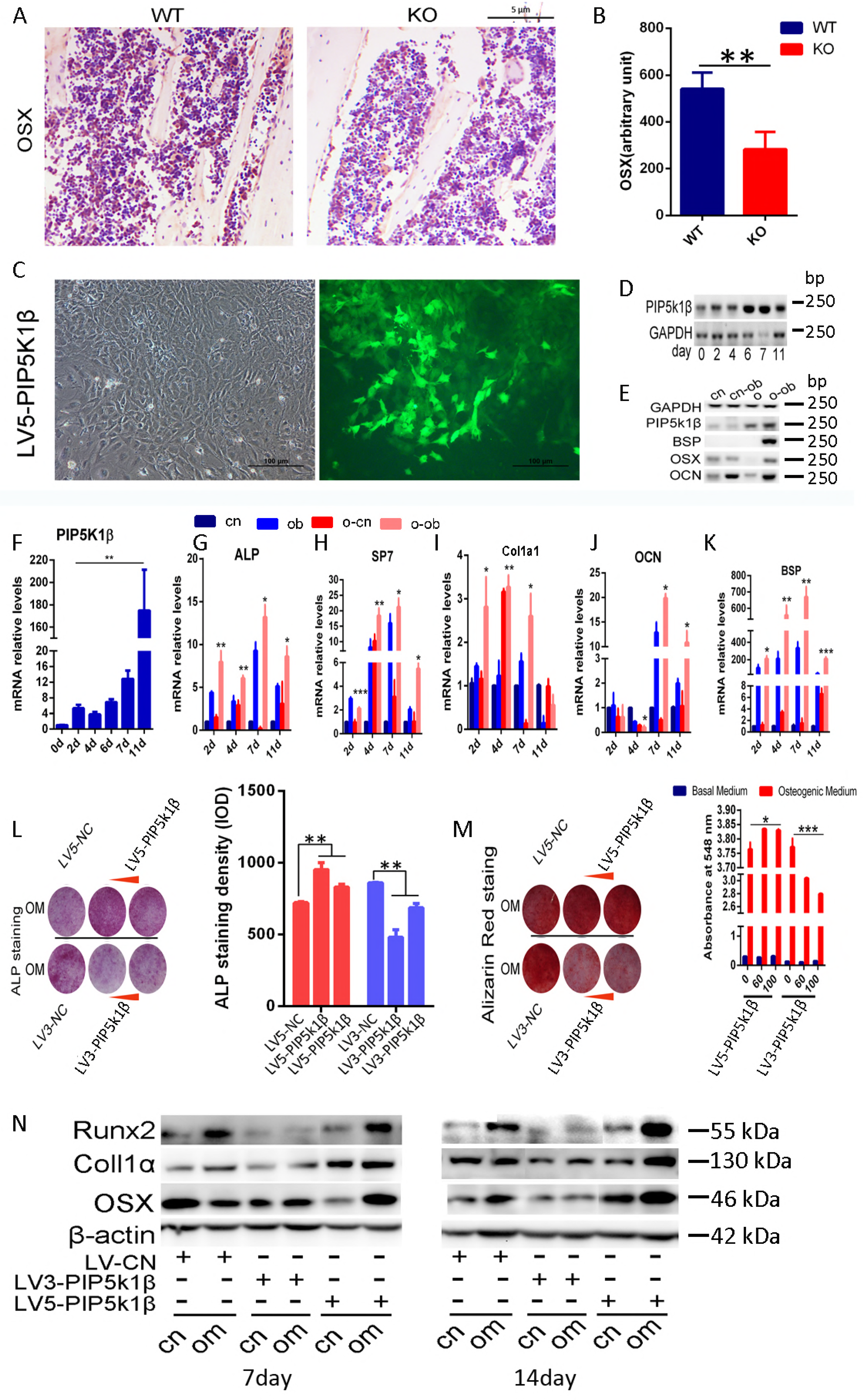
PIP5k1β accelerates osteoblast differentiation. (A-B) The expression of OSX in WT or PIP5k1β knockout mice shown by immunohistochemistry (A) and the staining density were quantified by ImageJ (B). (n=6 per group) (C) WT BMSCs were transfected with vectors or LV3-PIP5k1β (knockdown) or LV5-PIP5k1β (overexpression) to obtain BMSCs that constantly knockdown or overexpress PIP5k1β, the transfection ratio of LV5-PIP5k1β were shown. (D-K) BMSCs from C were cultured with osteogenic medium for the indicated times. RT-PCR was used to test PIP5k1β, BSP, OSX, OCN expression for an indicated number of times (D-E) and quantitative real-time PCR was performed for the mRNA expression of PIP5k1β, Alpl (ALP), Ibsp (BSP), SP7, OCN and Colla1 (F-K). (Data represent as means± SD of triplicate samples. cn, control cells treated with control medium; o and o-cn, cells overexpress PIP5k1β treated with control medium; cn-ob, control cells treated with osteogenic medium; o-ob, cells overexpress PIP5k1βtreated with osteogenic medium.) (L) Cells from C cultured for 7 days with osteogenic medium were fixed and stained for ALP. (M) Cells cultured with osteogenic medium for 14 days were fixed and stained for Alizarin red and were quantified by densitometry at 548 nm. (N) Western blot for osteoblasts marker genes for 7 and 14 days treatment with osteogenic medium and β-actin were used as loading control. (All experiments were performed triplicate. Data represent means ± SD of triplicate samples. *P < 0.05; ** P < 0.01; ***P < 0.001 versus control, cn, cells treated with control medium; om, cells treated with osteogenic medium.)

### PIP5k1β enhances osteoblast differentiation through activating p-Smad1/5/8 signaling

Several signaling pathways were involved in regulation of BMSCs differentiation into osteoblast, of which smad1/5/8 signaling exerts crucial roles in osteogenesis. To understand through which pathways PIP5k1β modulates osteoblast differentiation, we tested the activation of main pathways that regulates MSCs differentiation into osteoblast including smad1/5/8 and β-catenin and found that PIP5k1β prominently modulates smad1/5/8 activation. The phosphorylation of smad1/5/8 was significantly increased after 7 and 14 days treatment with osteogenic medium compared with cells treated with control medium, while, knockdown of PIP5k1β drastically depressed smad1/5/8 phosphorylation but overexpression of PIP5k1β outstandingly increased its phosphorylation on the contrary (Fig. 8A-B). To refine this finding, we further treated BMSCs with LDN193189, a smad1/5/8 specific inhibitor, to eliminate the activation of smad1/5/8. We observed that PIP5k1β overexpression prominently enhanced osteoblast differentiation and LDN193189 significantly reduced osteogenesis, but the reduction ratio was partly rescued by overexpression of PIP5k1β (Fig. 8C-E). Western blotting showed that PIP5k1β overexpression violently enhanced osteoblastogenesis master genes, Col1α1, Runx2 and OSX, expression, and LDN193189 significantly inhibited expression of these genes, but the inhibition ratio was partly rescued by overexpression of PIP5k1β. Meanwhile, Phosphorylation of smad1/5/8 was changed in consistent with the change trend of these osteoblastogenesis master genes. (Fig. 8F). In summary, these observations indicated that PIP5k1β enhances BMSCs differentiation into osteoblast partly through activating smad1/5/8 signaling.

**Figure 8.**
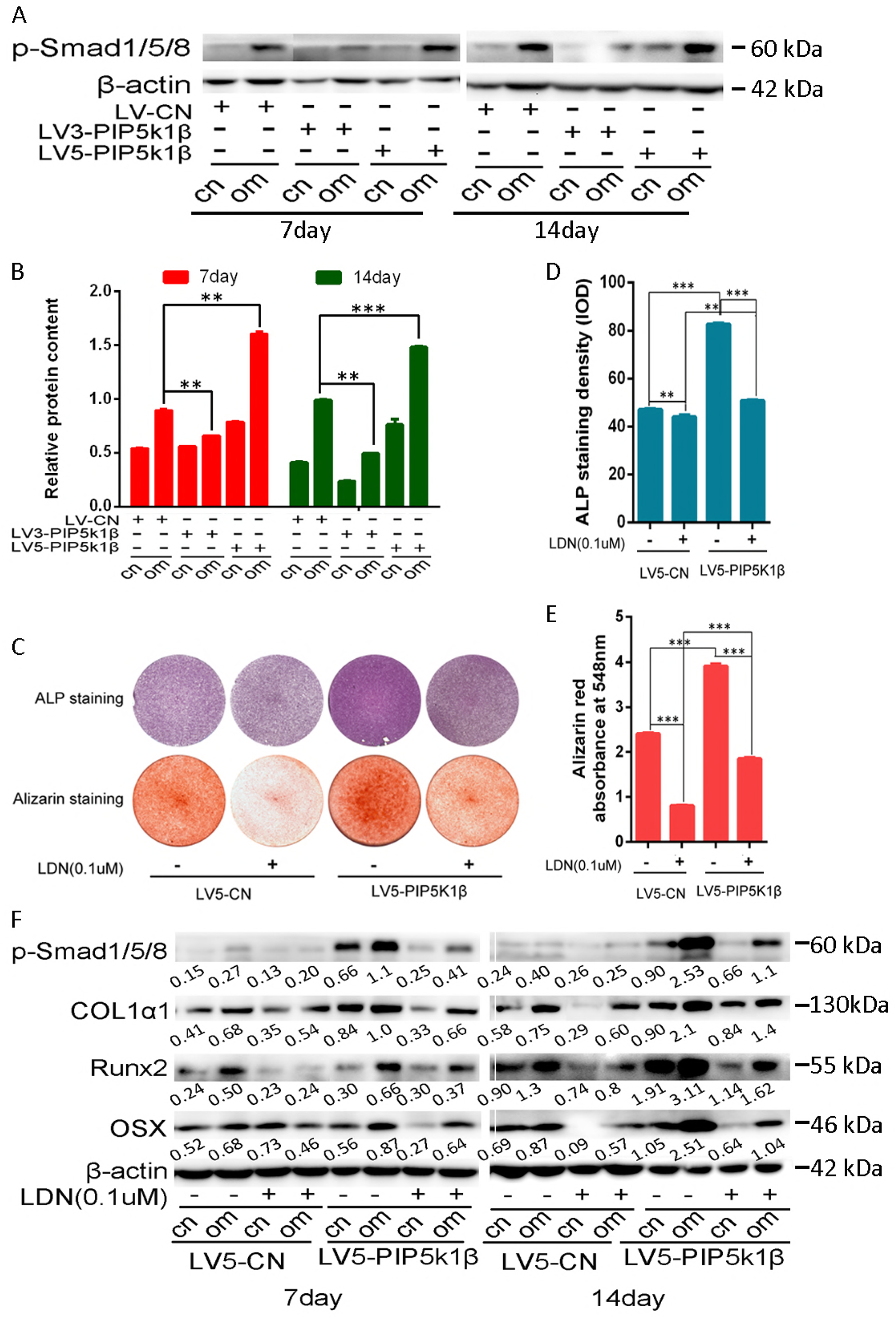
PIP5k1β accelerates osteoblast differentiation through activating p-Smad1/5/8 signaling. WT BMSCs were transfected with vectors or LV3-PIP5k1β (knockdown) or LV5-PIP5k1β (overexpression) to obtain BMSCs that constantly knockdown or overexpress PIP5k1β. Then were underwent with control or osteogenic medium incubation for 7 or 14 days. (A) Western blot for p-smad1/5/8 after 7 and 14 days treatment with osteogenic medium and β-actin were used as loading control. (B) The band densitometry from A was analyzed with ImageJ and was shown graphically. (C) ALP and Alizarin Red S staining for BMSCs incubated with control or osteogenic medium with or without smad1/5/8 specific inhibitor LDN193189 for 7 and 14 days respectively. (D) The ALP staining density was analyzed using the software Image-Pro Plus and shown as graph. (E) The calcium precipitates in figure B were dissolved in 0.1 N sodium hydroxide and quantified by absorbance at 548 nm with a Tecan Safire2 microplate reader (Tecan, Durham, NC, USA) (F) Westernblotting for BMSCs incubated with control or osteogenic medium with or without smad1/5/8 specific inhibitor LDN193189 for 7 and 14 days respectively. Expression of p-smad1/5/8 and osteogenic marker genes were detected and analyzed with ImageJ. (All experiments were performed triplicate. Data represent means ± SD of triplicate samples. *P < 0.05; ** P < 0.01; ***P < 0.001 versus control, cn, cells treated with control medium; om, cells treated with osteogenic medium.)

## Discussion

Several therapeutic strategies have been developed for excess bone loss diseases such as osteoporosis, including direct suppression of osteoclasts resorption capacity or irritation of osteoblast bone formation, with compromised efficacy and momentous side effects. Therefore, more appropriate treatments need to be explored to synchronously modulate the interaction between osteoblasts and osteoclasts. In present study, we discovered that PIP5K1β inhibited osteoclasts formation and function but meanwhile facilitated osteoblast differentiation, so that PIP5k1β^−/−^ displayed obvious osteoporosis phenotype. These hinted that PIP5K1β might serve pivotal roles in maintaining bone homeostasis and that deepening understanding of the mechanisms of its functional role might provide new strategy for the prevention and treatment of osteoporosis.

Among all the family members of PIP5k1 family kinases that generate PIP2, only PIP5k1β was highly expressed in bone and during osteoblastogenesis and RANKL-induced osteoclast differentiation, which indicated that PIP5k1β might serve crucial roles in bone biology. Under the consideration that PIP5k1 family proteins were reported to be involved in cytoskeleton assembly and that the podosome reorganization of osteoclasts was a major event for mature osteoclasts formation and activity(Mao & Yin, 2007; Teitelbaum, 2011; van den Bout & Divecha, 2009), we speculated that PIP5k1β might participate in osteoclast sealing zone organization. However, our present study showed that PIP5k1β deletion osteoclasts exhibit no apparent cytoskeletal abnormalities and also had no impact on the expression and colocalization of proteins involved in sealing zone formation, such as vinculin, Rac1 and F-actin. Additionally, the expression levels of β3 integrin, which takes part in the association of osteoclasts with bone matrix to trigger the osteoclast bone resorption activity, was also uninfluenced by PIP5k1β deficiency (Supplementary Fig. 1C, D). This accumulating evidence indicated that PIP5k1β affects osteoclasts not by affecting their cytoskeleton formation and interaction with bone matrix though deletion of PIP5k1β accelerates and enlarges osteoclast actin ring formation. Nonetheless, PIP5k1β indeed influences osteoclast formation and function, which might be buttressed by the following facts. First, PIP5k1β deletion enhanced the expression of osteoclasts marker genes such as ACP5, CTSK, DC-STAMP, c-Fos, NFATC1 and so on. Besides, PIP5k1β deficiency facilitated osteoclast bone resorption as proven by enhanced pit formation activity and highly elevated serum level of bone resorption marker CTX-1 in PIP5k1β deletion mice. Finally, knockdown and overexpression of PIP5k1β in BMMs accelerated and depressed osteoclast differentiation, respectively, as detected by TRAP staining.

Osteoclasts are generated from mononuclear hematopoietic myeloid lineage cells, which are derived in the bone marrow and are attracted to the blood stream by factors. These circulating precursors migrated to bone surfaces undergoing resorption by chemokines and other factors that were modulated by M-CSF and RANKL at these sites, where they fuse to develop multinucleated bone resorbing cells(Kikuta et al, 2011; Kular et al, 2012). Interestingly, in the present study, we obtained that PIP5k1β deletion can accelerate preosteoclast migration triggered by M-CSF or by both M-CSF and RANKL. Besides, PIP5k1β deletion promoted preosteoclast proliferation. These results indicated that PIP5k1β repressed preosteoclast proliferation and migration to inhibit osteoclastogenesis, but the molecular mechanisms remain to be explored.

Furthermore, our present study suggest that PIP5k1β affected osteoclast differentiation and function also through modulation of NFATC1 which represents a master switch that modulates terminal differentiation of osteoclasts downstream of RANKL(Asagiri et al, 2005; Takayanagi et al, 2002), as supported by several discoveries. First, we found that PIP5k1β deficiency enhanced NFATC1 expression and facilitated NFATC1 nuclear translocation. Additionally, overexpression of PIP5k1β dampened NFATC1 activity. Moreover, the phosphorylation of Akt and the activation of MAPK/p38, ERK1/2 and JNK, which mediated NFATC1 activation during RANKL-stimulated osteoclast differentiation, were prolonged and enhanced with RANKL stimulation in PIP5k1β^−/−^ osteoclasts. Last but not the least, PIP5k1β deletion enhanced TRAF6 and c-Fos expression, which were demonstrated to stimulate the downstream mediators to trigger expression and nuclear translocalization of NFATC1 resulting in enhanced osteoclastogenesis.(Boyce & Xing, 2008; Kadono et al, 2005) (Wang et al, 1992) PIP5K1γ was reported to modulate calcium transport, which in turn, initiated NFATc1 (Wang et al, 2004). Moreover, NFATC1 can autoamplify and trigger calcitonin receptor expression which eventually upregulated calcium signaling (Asagiri et al, 2005). Further studies need to be performed to determine if PIP5k1β deletion affects calcium signaling.

Adaptor protein Grb2 (growth-factor-receptor-bound protein 2), which serves an important role in several tyrosine kinase signal transduction pathways, was reported to play a crucial role in osteoclastogenesis by increasing Erk and Akt signaling motivated by M-CSF to promote osteoclast proliferation, differentiation and survival.(Levy-Apter et al, 2014; Ross & Teitelbaum, 2005; Takayanagi, 2007) An amusing result was discovered in the present study that Grb2 expression was advanced and dramatically up-regulated during osteoclastogenesis while PIP5k1β was deleted, besides, the ERK1/2 signaling was also enhanced. These implied that PIP5k1β might modulate Grb2 expression to regulate osteoclastogenesis. However, further study is needed to make deep insight in the association between PIP5k1β and Grb2 expression.

Our present work also discovered that PIP5k1β facilitated osteoblast differentiation, immunohistochemistry in Fig. 7A showed that OSX^+^ cells was prominently decreased both in bone marrow cells and in bone lining cells in PIP5k1β deletion mice compared with that of WT mice, which indicated that PIP5k1β might stimulate MSCs differentiation into osteoblast through up-regulation of OSX expression. Furthermore, the expression of osteoblast differentiation early and late markers Col1α1, ALP, OCN as well as BSP and osteoblast differentiation master transcription factor Runx2 and OSX were all outstandingly elevated or reduced during osteoblastogenesis of PIP5k1β overexpression or knockdown BMSCs respectively. Moreover, ALP and Alizarin Red staining demonstrated that overexpression or knockdown of PIP5k1β promoted or repressed MSCs differentiation into osteoblast and calcium precipitation, respectively. Meanwhile, the phosphorylation of smad1/5/8 was also enhanced or dampened in accordance with overexpression or knockdown of PIP5k1β during BMSCs differentiation into osteoblast. Moreover, PIP5k1β overexpression can partly rescue the elimination effect of osteoblast differentiation and smad1/5/8 activation upon LDN193189, a smad1/5/8 specific inhibitor, administration. These accumulating evidences implied that PIP5k1β facilitates osteoblast differentiation from BMSCs partly through smad1/5/8 signaling.

In summary, our study explored the role of PIP5k1β in bone biology, particularly in osteoclast and osteoblast differentiation and function, for the first time. We found that PIP5K1β was highly expressed both during osteoclast and osteoblast differentiation and PIP5K1β can regulate bone mass and bone remodeling by inhibiting osteoclast differentiation and facilitating osteoblast differentiation. (Supplementary Fig. 2D) Further studies need to be performed to determine if these functions were partly caused by PIP2 levels alteration that was contributed by PIP5k1β kinase activity during osteoclast and osteoblast differentiation. Besides, further explanation of the RANKL modulation of PIP5k1β expression is needed to deepen our understanding of the biological roles of PIP5k1β on maintaining bone homeostasis to explore new targets for prevention and treatment of bone disorders.

## Materials and Methods

### Reagents

Alpha-minimum essential medium (MEM), fetal bovine serum (FBS), and penicillin were purchased from Gibco BRL (Gaithersburg, MD, USA). Recombinant mouse macrophage-colony stimulating factor (M-CSF) was purchased from R&D Systems (USA). Recombinant Murine sRANK Ligand was purchased from Peprotech (USA).Tartrate-resistant acid phosphatase (TRAP) staining solution was obtained from Sigmae-Aldrich (St. Louis, MO, USA). The Cell Counting Kit-8 (CCK-8) was obtained from Dojindo Molecular Technology (Japan).

### Animals

FVB WT and PIP5k1β^−/−^ mice were obtained from PBmice of Fudan University (http://www.idmshanghai.cn/PBmice or http://www.scbit.org/PBmice/) and housed five per cage under standard conditions (12 h light/12 h dark cycle, 21°C controlled temperature).

### Sample preparation and skeletal morphology

For microCT analysis, right femurs of both WT and PIP5k1β^-^/^-^ mice were fixed with 4% paraformaldehyde and analyzed by Scanco Medical CT-40 instruments. Trabecular bone analysis was performed on the secondary spongiosa region (300 μm below the growth plate with a total height of 1 mm towards the mid shaft) of the distal femur. 3D images were generated in CTvol program (Skyscan). For histochemistry, proximal tibias from both WT and PIP5k1β^−/−^ mice were fixed in 4% paraformaldehyde and embedded in paraffin, then cut longitudinally into 4-mm-thick sections and processed for hematoxylin and eosin staining (H&E staining) or Tartrate-resistant acid phosphatase (TRAP) staining after decalcifying. To evaluate bone formation rate in vivo, WT and PIP5k1β^−/−^ mice were injected with green fluorescent calcein (Sigma-Aldrich; 5 mg per kg body weight) on 8 and 2 days before euthanasia. Then the tibiae were dissected and embedded in methyl methacrylate resins. Tissue sections were observed under a laser-scanning microscope (LSM5 PASCAL; Carl Zeiss)

### Cell Culture

WT or PIP5k1β^−/−^ BMMs were obtained as described previously. (Li et al, 2013; Qin et al, 2012)In brief, bone marrow cells extracted from the femurs and tibiae of a 9- to 10-week-old WT or its littermate PIP5k1β^−/−^ mouse were cultured in α-MEM medium containing 15 ng/ml M-CSF (416-ML; R&D; USA) in a T-75 cm^2^ flask for proliferation for 2–3 days until reaching 90% confluence, the cells were washed with PBS three times and trypsinized for 30 min to harvest BMMs. Adherent cells were classified as BMMs. WT and PIP5k1β^−/−^ BMMs were plated on 96-well plates at a density of 8×10^3^ cells/well in triplicate and incubated with α-MEM medium containing 20 ng/ml M-CSF in a humidified incubator containing 5% CO2 at 37°C for 2 d. Then, the cells were cultured with α-MEM medium with M-CSF (20 ng/mL) and RANKL (75 ng /mL; 315-11; Peprotech; Rocky Hill, NJ) for indicated times. Moreover, the medium was changed every other day. For osteoblast differentiation, bone mesenchymal stem cells (BMSCs) were gained from 9- to 10-week-old mice and expanded as previously described, induced by culturing cells in osteogenic medium (DMEM containing 1M β-glycerophosphate, 50mM ascorbic acid and 1mM Dex (methylisobutylxanthine)) for indicated times, followed by subsequent experiments.

### TRAP activity assay

After 7 days of culture, the osteoclasts were fixed with 4% paraformaldehyde (PFA) in PBS for 10 min and then rinsed three times with PBS, followed by TRAP staining, using an acid phosphatase kit (387A; Sigmae^_^Aldrich) according to the manufacturer’s description and counter-stained with hematoxylin for 10 s to 30 s. Images were obtained with a Nikon SMZ 1500 stereoscopic zoom microscope (Nikon Instruments Inc.; Melville, NY, USA). We randomly quantified the total area of TRAP-positive regions and the total number of osteoclasts on five selected fields of view for each sample.

### Immunofluorescence

For F-actin ring immunofluorescent staining, WT and PIP5k1β^−/−^ osteoclasts were fixed with 4% PFA for 15 min at room temperature and permeabilized for 5 min with 0.1% v/v Triton X-100 on ice. Cells were incubated with Fluorescent phalloidins (1:40; Invitrogen Life Technologies, USA) diluted in 1% w/v bovine serum albumin-PBS for 20 min at room temperature and then washed extensively with PBS. Cells were then incubated with Hoechst 3342 dye (1:5000); Invitrogen Life Technologies, USA) for visualizing nuclei, washed with PBS, and mounted with ProLong Gold anti-fade mounting medium (Invitrogen Life Technologies, USA). Fluorescence was detected with NIKON A1Si spectral detector confocal system equipped with 20 (dry) lenses. Fluorescence images were obtained with NISeC Elements software. (National Institutes of Health).

### Pit formation assay

For the bone resorption assay, 8×10^3^ cells/well WT or PIP5k1β^−/−^ BMMs were plated onto bovine bone slices in 96-well plates with three replicates and treated with M-CSF (20 ng/mL) and RANKL (75 ng/mL) for 8 days. Then the bone slices were fixed with 2.5% glutaraldehyde and followed by visualization with scanning electron microscope (SEM; FEI Quanta 250). Pit areas were quantified with Image J software (National Institutes of Health). Similar independent experiments were repeated three times.

### Cell proliferation and migration assay

The proliferation rate was determined with CCK-8 assay according to the manufacturer’s protocols. WT or PIP5k1β^−/−^ BMMs were seeded in 96-well plates at a density of 8×10^3^ cells/well, and incubated in complete a-MEM supplemented with 20 ng/mL M-CSF for 24 h until 60% confluences, then cells were treated with 20 ng/ml M-CSF or 75 ng/ml RANKL or both for 24 h. Then 100 ul CCK-8 buffer was added to each well, followed by incubation at 37 °C for an additional 2 h. The absorbance was then measured at a wavelength of 450 nm (650 nm reference) with an ELX800 absorbance microplate reader (Bio-Tek, USA). The migration ability was monitored with wound healing assay. In a typical procedure, a straight scratch was made gently through the monolayer cells using an Eppendorf tip in 48-well plates. Detached cells were washed away with PBS and the serum-free culture medium was added; cells were treated with 20 ng/ml M-CSF or 75 ng/ml RANKL or both. Cells migrating to the scratch were monitored and images were taken at 0, 17h after wounding. The migration rate was calculated following the equation: percentage wound healing = ((wound length at 0 h) − (wound length at 17 h))/(wound length at 0 h) × 100 (Davalos et al, 2012).

### RNA extraction and Q-PCR assay

WT and PIP5k1β^−/−^ BMMs were plated in six-well plates at a density of 1×10^5^ cells per well and incubated in complete a-MEM supplemented with 20 ng/mL M-CSF and 75 ng/mL RANKL for an indicated number of days. Total RNA was extracted with TRIzol reagent (Invitrogen) following the manufacturer’s instructions and cDNA was generated with 1 ug of RNA. Then, real-time PCR was performed via the LightCycler480 system (Roche) using SYBR1Premix Ex TaqTM (Takara, Dalian, China) following manufacturer’s instructions and data were analyzed using the comparison Ct (2^-ΔΔ^Ct) method The following program of real-time PCR was employed: denaturation at 95 °C for 10 s, 40 cycles at 95 °C for 10 s, and 60 °C for 30 s. The dissociation stage was added to the end of the amplification procedure, and the dissociation curve did not show any nonspecific amplification. Glyceraldehyde-3-phosphate dehydrogenase (GAPDH) was used as housekeeping gene. All reactions were run in triplicate. The mouse primers used in the present study were shown as follows: GAPDH forward 5’-TTCACCACCATGGAGAAGGC-3’ and reverse 5’-GGCATGGACTGTGGTCATGA-3’; Nfatc1 forward 5’-TCCGAGAATCGAGATCACCT-3’ and reverse 5’-AGGGGTCTCTGTAGGCTTCC-3’; Acp5 (acid phosphatase 5) forward 5’-CGTCTCTGCACAGATTGCAT-3’ and reverse 5’-AACTGCTTTTTGAGCCAGGA-3’; cathepsin K (CTSK) forward 5’-GGACCCATCTCTGTGTCCAT-3’ and reverse 5’-CCGAGCCAAGAGAGCATATC-3’; DC-STAMP _dendritic cell-specific transmembrane protein (DC-STAMP) forward 5’-ACAAACAGTTCCAAAGCTTGC-3’ and reverse 5’-TCCTTGGGTTCCTTGCTTC-3’; OSCAR forward 5’-CTGCTGGTAACGGATCAGCTCCCCAGA-3’ and reverse 5’-CCAAGGAGCCAGAACCTTCGAAACT-3’; PIP5k1β forward 5’-GAAGAAGCCCTGGGATCCCGACA-3’ and reverse 5’-GGGTGTTTGGCTCAGCCGTCA-3’; ATP6v0d2 forward 5’-AGACCACGGACTATGGCAAC-3’ and reverse 5’-CAGTGGGTGACACTTGGCTA-3’

### Western blotting

Cells were lysed on ice for 30 min with RIPA lysis buffer which contains 50 mM Tris^_^HCl, pH 7.4, 150 mM NaCl, 1% Nonidet P-40, and 0.1% SDS supplemented with protease inhibitors (10 mg/ml leupeptin, 10 mg/ml pepstatin A, and 10 mg/ml aprotinin). For Western blotting, 25µg of protein sample was resolved on 12.5% SDS-PAGE and electrotransferred onto nitrocellulose membranes (Whatman, Piscataway, NJ, USA). The primary antibodies used were as follows: phospho-Akt, total Akt, phosphor –GSK3β(ser9A), PTEN, PLCγ2 and all the primary antibodies involved in the MAPK signaling pathway (phospo-p44/42 ERK, total p44/42 ERK, phospho-p38, total-p38, and phospo-JNK and total-JNK) were purchased from Cell Signaling Technology (Danvers, MA, USA) and used at a 1:1000 dilution ratio. Anti-NFATC1 antibody was purchased from Santa Cruz and used at 1:500 dilution ratio. Beta-actin or GAPDH was used as loading control. HRP-conjugated secondary antibodies were used at a 1:5000 dilution. The antigen–antibody complexes were visualized using the enhanced chemiluminescence detection system (Millipore, Billerica, MA, USA) following the manufacturer’s instructions. Immunoreactive bands were quantitatively analyzed in triplicate by normalizing the band intensities to their respective controls on scanned films using ImageJ software.

### Alkaline phosphatase staining and Alizarin red staining

The cell layer was rinsed with PBS three times, followed by fixation in 4% paraformaldehyde for 10 min at room temperature. Then cells underwent ALP or Alizarin red staining. ALP staining was performed according to the manufacturer’s instructions. The fixed cells were then incubated with buffer containing 0.1% naphthol AS-Bi phosphate (Sigma^_^Aldrich) and 2% fast violet B (Sigma^_^Aldrich). After incubation for 1 h at 37 °C, the cell layer was washed with deionized water. For Alizarin red staining, the fixed cells were rinsed with double-distilled H2O (ddH2O) and then stained with 40 mM Alizarin red S (pH 4.9, Sigma) for 15 min with gentle agitation followed by five times washing with ddH2O.Then, the calcium precipitates were dissolved with 0.1 N sodium hydroxide and monitored using a Tecan Safire2 microplate reader (Tecan, Durham, NC, USA) by absorbance at 548 nm to quantify the degree of mineralization.

### Statistical analysis

The data are presented as the mean ± s.d. (n is the number of tissue preparations, cells, or experimental replicates). For comparing groups of data, a two-tailed Student’s *t*-test was used. A value of P < 0.05 was considered to be statistically significant.

## Author contributions

Xiaoying Zhao: Conception and design, manuscript writing, collection of data, data analysis and interpretation; Guoli Hu and Chuandong Wang: Collection of data, Provision of study material; Lei Jiang and Jingyu Zhao: Collection of data and manuscript revision; Jiake Xu: Data analysis and manuscript revision; Xiaoling Zhang: Conception and design, manuscript revision, final approval of manuscript.

## Acknowledgments

This work was supported by grants from National Natural Science Foundation of China (No.81572123, 81401844), Science and Technology Commission of Shanghai Municipality (No.14431900900, 15411951100, 16430723500), Shanghai Municipal Education Commission-Gaofeng Clinical Medicine Grant Support (No.20161314) and Xin Hua Hospital Affilliated to Shanghai Jiao Tong University School of Medicine (009&201601).

---

**Supplementary Figure 1.** (A) The whole body of WT and PIP5k1β deletion mice. (B) Total RNA was isolated from bone, spleen, brain, thymus, lung, kidney, liver, epididymis, testis, heart, and muscle of WT mice. Quantitative real-time PCR was performed for the mRNA expression of PIP5k1α, β and γ. (C-D) WT or PIP5k1β−/− BMMs were cultured with M-CSF (20 ng/mL) and RANKL (75 ng/mL) for 0, 1, 3, 5 or 7 days. β3 Integrin expression was analyzed by real-time polymerase chain reaction (PCR) and the results were normalized to the expression of GAPDH (C) and western blotting (D). All experiments were performed at least three times (*P < 0.05; **P < 0.01)

**Supplementary Figure 2.** (A) MTT assay to detect the proliferation rate of BMSCs constantly overexpress or knockdown of PIP5k1β and the control BMSCs. (B-C) Densitometry analysis of Fig.7N and the results were shown graphically. (All experiments were performed triplicate. Data represent means ± SD of triplicate samples. *P < 0.05; ** P < 0.01; ***P < 0.001 versus control, cn, cells treated with control medium; om, cells treated with osteogenic medium.) (D) Model of PIP5k1β to modulate bone homeostasis. On the one hand, RANKL binds to RANK then RANK recruit TRAF6, which leads to phosphorylation of JNK, Akt MAPK/Erk1/2 and MAPK/p38 signaling and activation of c-Fos, which are crucial for NFATC1 induction and activation. NFATC1 in turn initiates its own expression and activation. NFATC1 promotes osteoclastogenesis by promoting expression of osteoclastogenic genes such as DC-STAMP, Cathepsin k, OSCAR, ACP5, Atpasev0d2. PIP5k1β inhibits RANKL-induced MAPK, AKT activation and TRAF6, c-Fos expression; PIP5k1β also inhibits Grb2 expression during osteoclastogenesis. Grb2 promotes osteoclasts proliferation and formation. In this way, PIP5k1β inhibits osteoclastogenesis to modulate bone resorption. On the other hand, PIP5k1β promotes expression of osteogenic genes such as ALP, OSX, RUNX2, OCN, BSP, col1α1 to enhance bone formation through activating smad1/5/8 signaling.

